# A Life Identification Number Barcoding (LIN Code) System for *Neisseria meningitidis*: high resolution multi-level typing of meningococci

**DOI:** 10.64898/2026.03.03.708563

**Authors:** Kasia M Parfitt, Keith A Jolley, Anastasia Unitt, James E Bray, Frances M Colles, Odile B Harrison, Ian M Feavers, Martin C J Maiden

## Abstract

*Neisseria meningitidis* is a commensal member of the human oropharyngeal microbiota that can cause devastating invasive meningococcal disease. This genetically and antigenically diverse ‘accidental pathogen’ has been a paradigm for the study of bacterial population biology. The meningococcus was the first organism for which a seven-locus multi-locus sequence typing (MLST) scheme was developed. With the addition of sequence-based characterisation of antigen genes, molecular typing has been widely employed to inform surveillance and public health interventions. Following the advent and widespread adoption of whole genome sequencing (WGS), precise delineation of variants is possible. Here, a WGS-based Life Identification Number (LIN) code typing scheme is described, providing a multi-resolution nomenclature for understanding meningococcal population diversity and molecular epidemiology. The LIN codes were developed using a set of 6,131 *N. meningitidis genomes*, comprising up to 200 isolates from each clonal complex (cc) previously described using MLST. Based on cluster-stability analysis and concordance with ccs, thirteen LIN ‘thresholds’ described the meningococcal population at different levels of resolution. LIN codes and human-readable ‘nicknames’ consistent with existent nomenclatures were assigned to the *N. meningitidis* genomes hosted in the PubMLST. Published outbreaks validated the LIN thresholds, illustrating the potential of this genomic tool in public health management.

## Introduction

*Neisseria meningitidis*, the meningococcus, is a common member of the human oropharyngeal microbiota that occasionally causes devastating invasive meningococcal disease (IMD) (Tzeng and Stephens, 2021; Pardo de Santayana *et al*., 2023). IMD occurs in multiple clinical and epidemiological manifestations including endemic disease, outbreaks at various scales (Jolley *et al*., 2012), and large-scale epidemics and pandemics (Mbaeyi *et al*., 2020). Predominantly presenting as meningitis and or septicaemia, with a case fatality rate of ranging from 5-16% (Cabellos *et al*., 2019), IMD includes other syndromes such as pneumonia, chronic arthritis, and pericarditis (Stephens *et al*., 2007). IMD presentation and epidemiology is strongly influenced by the genetic and antigenic diversity of this highly recombinogenic bacterium. There are a limited number of genotype:antigen type combinations dominating IMD isolate collections, and greater diversity is present in isolates from asymptomatic carriers than IMD (Stollenwerk *et al*., 2004; Caugant and Maiden, 2009).

The first characterisation methods of the meningococcus were phenotypic, including serological methods. Increasingly from the 1990s, molecular characterisation of meningococcal variants (Caugant and Nicolas, 2007) played a central role in improvements in global control of IMD (Jolley *et al*., 2007; Slavinska *et al*., 2024). These molecular methods have the advantages of being rapid, exact, and readily reproducible. They can also identify hyper-invasive meningococcal lineages: related genotypes especially associated with IMD, enabling improved understanding of epidemiology (Jolley *et al*., 2012; Hamed *et al*., 2021), pathology, antimicrobial resistance, and disease prevention (Guo *et al*., 2018; Chen *et al*., 2021). Multi-locus sequence typing (MLST) leveraged developments in nucleotide sequencing technologies to index sequence variation in housekeeping genes, replacing Multi-Locus Enzyme Electrophoresis (MLEE) (Caugant *et al*., 1986; Reeves *et al*., 1995). This provided improved resolution, reproducibility, and portability through the distribution of such nomenclatures on the World Wide Web (www) via PubMLST.org (Caugant and Maiden, 2009; Maiden *et al*., 2013). This enabled the assignment of variants to sequence types (STs) and clonal complexes (ccs), as surrogates for genetic lineage (Jolley *et al*., 2007; Retchless *et al*., 2021). Seven-locus MLST has a wide range of public health and research applications including outbreak studies and long-term epidemiological study (Feavers *et al*., 1999; Birtles *et al*., 2005), particularly when supplemented with characterisation of fine-typing antigen genes (such as those encoding FetA and PorA). However, there are cases when highly similar meningococcal variants, not distinguished by MLST and fine typing, have distinct epidemiological properties. An example being the ‘Hajj’ and ‘South American/UK’ variants of the W:cc11 complex, which were discriminated by *ad hoc* whole genome sequences (WGS) analyses (Taha *et al*., 2000; Hahné *et al*., 2002; Lingappa *et al*., 2003) that included the first use of genomics to influence changes to a national immunisation programme to vaccinate individuals against meningococcal serogroups A, C, W(135), and Y (Hahné *et al*., 2002; Lucidarme *et al*., 2015; Badur *et al*., 2022; Campbell *et al*., 2022). Despite such successes, there remained a lack of a high-resolution, stable, and disseminated genome-level schemes to support public health interventions against the meningococcus over time and geographic spread. Core genome MLST (cgMLST) forms the basis of such approaches, but at the time of writing this was not systematised into a stable nomenclature schema.

Life Identification Number (LIN)® (Trademark registered by This Genomic Life, Inc, Floyd, VA, USA) codes exploit and catalogue variation present in cgMLST data through a multi-position, integer-based barcode which is automatically assigned to each isolate and describes its clustering at hierarchical thresholds of cgMLST allelic mismatch. LIN codes can be used at scale to define different levels within the population structure based on these thresholds (Jansen van Rensburg *et al*., 2024; Palma *et al*., 2024), providing an increased resolution compared to seven locus MLST. LIN code thresholds are chosen pragmatically, based on discontinuities in the population structure of the subject organism. Initially, the LIN concept was applied to levels of nucleotide sequence identity (Vinatzer *et al*., 2017), but here it is used on allelic comparisons within a cgMLST scheme (Hennart *et al*., 2022; Palma *et al*., 2024). LIN codes are assigned to each isolate based on its genomic similarity to other previously assigned isolates. As these are fixed, they are unaffected by the addition of further genomes into the scheme, providing stability to the nomenclature (Hennart *et al*., 2022). The principal application of LIN codes as a computer-readable numeric code, but accessibility to humans can be achieved through the assignment of ‘nicknames’ that are consistent with existing nomenclatures at the appropriate threshold (Palma *et al*., 2024). At the time of writing, LIN code schemes have been created for several bacterial pathogens including *Klebsiella pneumoniae* (Hennart *et al*., 2022), *Streptococcus pneumoniae* (Jansen van Rensburg *et al*., 2024), *Neisseria gonorrhoeae* (Unitt *et al*., 2025), and *Corynebacterium diphtheriae* (Delgado-Blas *et al*., 2025).

The meningococcal LIN code scheme described here was based on an updated cgMLST scheme (version 3) consisting of 1,329 loci and over 72,000 *N. meningitidis* genomes in PubMLST as of March 2025 (Jolley *et al*., 2018), and validated with previously published datasets. The LIN codes were then implemented and made publicly available in the PubMLST database (Jolley *et al*., 2018), providing an accessible and standardised multi-level *N. meningitidis* nomenclature.

## Methods

### Updating of the *Neisseria meningitidis* Core Genome MLST scheme (cgMLST v3)

The cgMLST v3 scheme for *N. meningitidis*, available within the PubMLST platform (Jolley *et al*., 2018), was based on cgMLST v2 (now archived within PubMLST). Annotation issues were reviewed during version updates. Inconsistencies in automated locus annotation included: (i) loci absent from a small subset of genomes that are excluded as not core genes; (ii) loci with inconsistent start sites, often resulting from the assignment of alleles from different *Neisseria* species; (iii) loci with multiple allele assignments, due to either multiple copies within the genome or closely related paralogues. Where multiple alleles were being assigned for a locus in some assemblies, all previously defined alleles were aligned using CLUSTALW and manually reviewed for inconsistent start sites. Further checks were made using the BIGSdb GenePresence tool to identify loci that were completely absent from many genome assemblies and therefore could not be defined as core. Finally, all loci that were assigned in <98% of high-quality draft assemblies (defined as >2Mbp total length, <= 200 contigs, and 0 gaps) underwent successive rounds of manual curation to improve assignment levels. Loci that were still assigned in <98% assemblies were then removed from the scheme.

### Genome data for LIN code development

All publicly available *N. meningitidis* isolates in the PubMLST database (Jolley *et al*., 2018) (>39,000 on 2^nd^ August 2024) were filtered for quality. High quality draft genomes were defined for the purpose of this study as having ≤500 contigs and ≥99% allelic designation. A LIN code development dataset of 6,131 *N. meningitidis* isolates (Dataset 1) (Supplementary Table 1) were chosen at random using the dplyr (v1.1.4) (Wickham, 2007) package in RStudio (v4.3.1) (RStudio Team, 2020). Up to 200 isolates from each clonal complex were used. MSTClust (v0.21b) (https://gitlab.pasteur.fr/GIPhy/MSTclust) (Hennart *et al*., 2022) was used to generate a pairwise distance matrix comparing cgMLST v3 across the 6,131 isolates (Dataset 1).

Reshape2 (v1.4.4) (Wickham, 2007) was used to melt the pairwise distance matrices to long format in Rstudio (v4.3.1) (RStudio Team, 2020). Ridgeline plots were created using the ggridges package (v0.5.6) from ggplot2 (v3.5.1) (Wickham, 2016). These plots were used to visualise peaks and troughs (discontinuity) in the datasets, comparing the number of isolates to percentage (%) allelic mismatches.

### Statistical analysis of pairwise allelic mismatches

MSTClust (v0.21b) (Hennart *et al*., 2022) was used with default parameters to calculate the Silhouette index and adjusted Wallace coefficients. The Silhouette index measured the clustering consistency within the datasets, while adjusted Wallace coefficient was used to indicate agreement within the groups of clusters. All pairwise allelic mismatches for Dataset 1 (*n* = 6,131) were subject to both statistical analysis methods. The adjusted Rand Index (Hubert and Arabie, 1985), was used to indicate the similarity between clustering by cc, ribosomal ST (rMLST), and ST and each was compared against each LIN threshold for the development Dataset 1 (*n* = 6,131). This used the adjustedRandIndex function from the mclust package (v1.0.4) (Scrucca *et al*., 2023) in RStudio (v4.3.1) (RStudio Team, 2020).

### Phylogenetic analyses

A second dataset consisting of 1,888 isolates (Dataset 2) (Supplementary Table 2) was used for phylogenetic analyses and LIN code validation. Isolates were chosen using dplyr (v1.1.4), with up to 50 isolates from each cc selected. The Genome Comparator plugin on PubMLST was used to create a cgMLST (v3) nucleotide alignment of all loci (*n* = 1,329 loci) using MUSCLE. RAxML (v8.2.12) (Stamatakis, 2006) was used to generate a maximum likelihood tree using the GTRGAMMA model, supported by 100 bootstraps. ClonalFrame (v1.12) (Didelot and Wilson, 2015) was used to reconstruct the RAxML tree and adjust for recombination. The phylogenetic tree was visualised using ggtree (v3.9.1) (Yu *et al*., 2017) from the ggplot2 package (v3.5.1) (Wickham, 2016) in RStudio (v4.3.1) (RStudio Team, 2020).

Four validation datasets (Table 1) were chosen to further represent the *N. meningtidis* population structure, outbreak, and carriage scenarios. Phylogenetic trees were generated for the validation datasets using RAxML (v8.2.12) (Stamatakis, 2006) and ClonalFrame (v1.12) (Didelot and Wilson, 2015) as described previously. Figtree (v1.4.4) (Rambaut, 2018) was used to root phylogenetic trees based on a selected outgroup. Phylogenies were then annotated using a combination of ggplot2 (v3.5.1) (Wickham, 2016) in RStudio (v4.3.1) (RStudio Team, 2020) and Inkscape (v1.3.2) (Inkscape Team, 2023).

**Table 1.** Representative datasets used for the validation analysis of the LIN codes.

| Dataset | No. isolates | Reference |
| --- | --- | --- |
| MLST dataset | 107 | Maiden *et al.* 1998 |
| Serogroup A epidemic of cc-5 | 23 | Diallo *et al.* 2017 |
| University of Southampton outbreak | 12 | Jolley *et al.* 2012 |
| Czech Republic carriage study (cc-11:C) | 32 | Jolley *et al.* 2000 |

## Results

### Creation of a cgMLST (v3) schema

The *N. meningitidis* cgMLST scheme was updated to ensure that all loci could be automatically annotated upon data deposition into PubMLST. The threshold of missing loci allowed for isolates to be assigned the same cgMLST was decreased from 50 (v1) (Bratcher *et al*., 2014) to 25 (v2), which was retained in v3. A total of 93 loci included in v2 were removed in the v3 update (Supplementary Table 3). This included: 8 removed due to unresolvable inconsistent start sites following assignment of alleles in different *Neisseria* species within the database; 15 removed as they were present in more than one copy in at least a subset of isolates; and 70 removed as they were assignable in <98% of high-quality assemblies because either they were absent, or a subset of alleles contained internal stop codons. The total number of loci for the updated cgMLST v3 schema was 1,329.

### Defining allelic mismatch thresholds for the *N. meningitidis* population

Dataset 1 (*n* = 6,131) was used to identify discontinuities in the meningococcal population. The total genome lengths ranged from 2.013 Mbp to 2.463 Mbp (mean length 2.146 Mbp) with the number of contigs ranging from 1 to 500, with a mean of 166 (Supplementary Figure 1). The log_10_ N50 values were between 3.97 and 6.36, with an average of 4.67 and GC content ranged from 50.15% to 52.2% (mean = 51.68%) (Supplementary Figure 1). Pairwise allelic mismatches across the cgMLST v3 core genes showed that the overall population structure of Dataset 1 (*n* = 6,131) was negatively skewed (Figure 1A). Most pairs had a high frequency of allelic mismatches, whereas fewer pairs were genetically similar, reflecting the diversity of *N. meningitidis*. The greatest frequency of paired allelic mismatches across the population was 1,217/1,329 (91.67%). Overall, Dataset 1 shared at least 41 loci (3.11% allelic similarity) as there were no paired mismatches exceeding 1,288 loci (96.89%) (Figure 1A).

**Figure 1.**
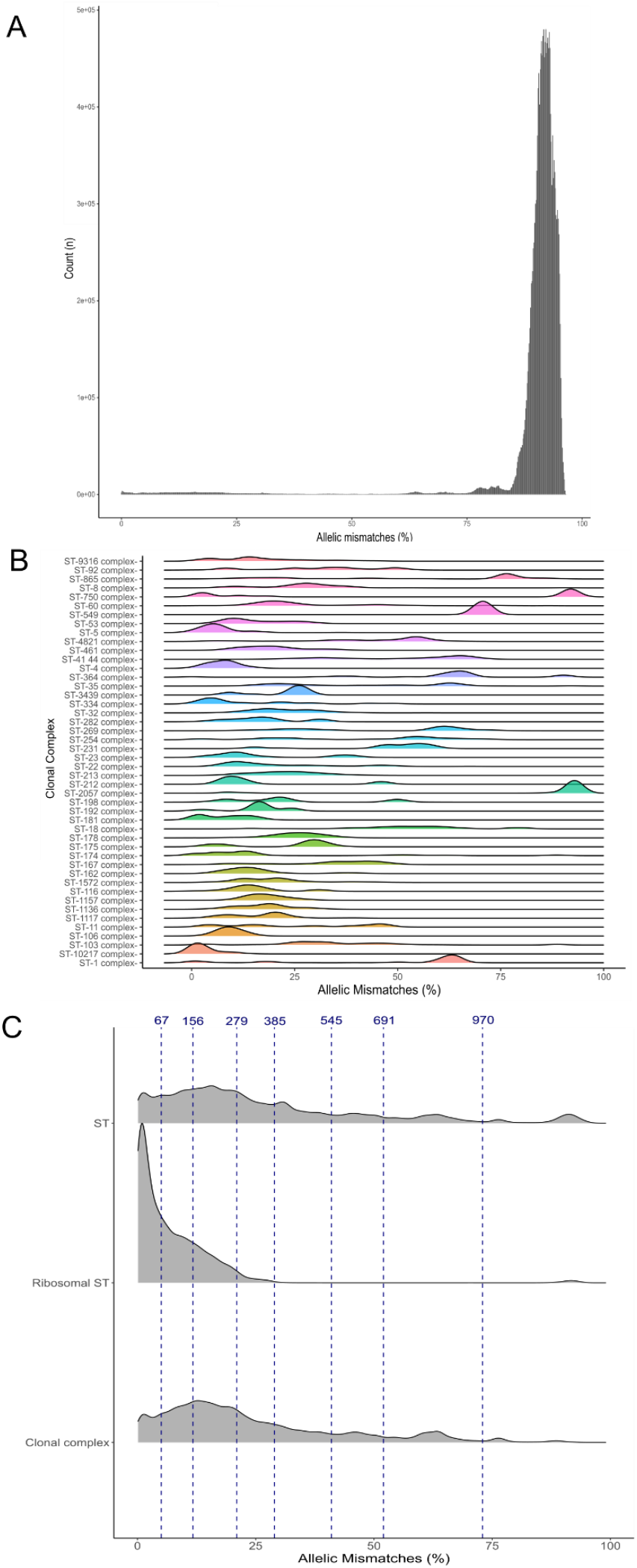
(A) Histogram showing the frequency (counts) of the percentage pairwise allelic mismatches (%) (n = 1,329 loci) within the N. meningitidis dataset (n = 6,131). This used an all-vs-all approach, using MSTClust (reference) to produce a distance matrix. (B) Ridgelines plot illustrating the frequency of allelic mismatches (%) within each cc (n = 6,129). The peaks illustrate the similarity of each cc population. ST-292 complex was removed from the analysis as there was only one comparison within the dataset, so a ridgeline could not be drawn. (C) Ridgeline plots showing the frequency of allelic mismatches for each matching metric: (i) ST, (ii) Ribosomal ST, and (iii) Clonal complex for the N. meningitidis isolates (n = 6,131). A subset of the allelic mismatch (%) thresholds are shown using blue dotted lines. These represent 72.96%, 52.00%, 41.00%, 28.97%, 21.00%, 11.74%, and 5.04% allelic mismatches (970, 691, 545, 385, 279, 156, and 67 loci mismatches, respectively).

There were clearer discontinuities within each cc, compared to the whole *N. meningitidis* population (Figure 1B). The distribution of allelic mismatches within each cc was variable. In some instances, there was only one major subgroup within that cc, e.g. within cc9316 (*n* = 121), cc549 (*n* = 4), cc5 (*n* = 199), cc106 (*n* = 15), and cc269 (*n* = 199). Within other clonal complexes, multiple subgroups were observed, including in cc1 (*n* = 110), cc865 (*n* = 154), cc4240/6688 (*n* = 4), and cc41/44 (*n* = 199), indicative of diversity and sub-lineages within some ccs. Most cc peaks were below 532 locus differences (40% allelic mismatches). The highest number of allelic mismatches across all ccs was observed in cc2057 at 93.37%. In contrast, cc4 had the fewest allelic mismatches (*n* = 9.48%) (Figure 1B; Supplementary Table 4).

To visualise the population structure of Dataset 1 in detail, pairwise comparisons were performed and compiled for all matching ccs, STs, and ribosomal STs (Figure 1B) This meant that a comparison was only created when both isolates belonged to the same clonal complex, which enabled identification of consistent subgroups across all metrics. All metrics showed the pairwise comparisons of the population to be of positive distribution (skewed to the left), and the number of allelic mismatches was less than 50% in most cases (Figure 1C).

An allelic mismatch threshold of 970 locus differences (72.96%) was chosen to represent ‘super-lineage’, which encompassed 99.22% of rSTs, 94.71% of MLSTs, and 97.87% of ccs. Cut-off values between 70.85-75.85% had corresponding Silhouette scores between 0.94-0.99 and adjusted Wallace scores of 0.87-0.98. A ‘lineage’ boundary at 691 locus differences (52% allelic mismatches) was chosen to correspond to cc. The corresponding Silhouette and adjusted Wallace scores were 1.00 and 0.95, respectively, indicating strong correspondence. This threshold captured 99.21% rSTs, 86.22% MLSTs, and 88.83% ccs. The next visible discontinuity below 665 locus differences (50% allelic mismatches) was consistent within both the ST and ccs metrics (Figure 1C). Around 80% of ccs and STs had 41% (*n* = 545 loci) or fewer allelic mismatches. Therefore, a threshold boundary was chosen at this level and named ‘sub-lineage’ (Silhouette score = 1.00 and adjusted Wallace score = 0.98).

Ribosomal MLST (rMLST) can be used for the rapid identification of bacterial species, lineage, and variants (Jolley *et al*., 2012). The majority of pairwise comparisons with matching ribosomal sequence types (rSTs) in Dataset 1 shared at least 385 loci (99.12% of *N. meningitidis* with the same rST and 70.33% ccs had less than 28.97% allelic mismatches). This threshold marked the end of the major rST tail. A similar observation was made for the 279 loci (21.00% allelic mismatch) boundary. This fell just before the rST tail (96.50% rSTs had less than 279 locus differences), cutting through the end of the major MLST and cc peak. The thresholds at 156 loci (11.74% allelic mismatches) and 67 loci (5.04% allelic mismatches) were chosen as they represented the median for all pairwise MLST and rST allelic mismatches, respectively. A total of 5 additional thresholds were defined with allelic mismatches of: 14, 7, 3, 2, 1 loci (corresponding to 1.05%, 0.53%, 0.23%, 0.15%, and 0.08% allelic mismatches respectively), to provide high discrimination for applications such as such as outbreak investigations. These were termed ‘clonal groups’. Indistinguishable isolates were those with identical matching LIN codes (0 locus differences; 0.00% allelic mismatches).

Before implementation on PubMLST, a local LINcoding Python script (https://gitlab.pasteur.fr/BEBP/LINcoding) was used to assign cgMLST-based LIN codes to each isolate in Dataset 1 (*n* = 6,131). To validate chosen thresholds, the adjusted Rand Index (ARI) (Hubert and Arabie, 1985) was used to infer concordance between LIN code clustering and pre-defined metrics. Concordance against cc, ST, and rST was assessed. At the 52% lineage allelic mismatch threshold (691 locus differences), there was strong concordance between LIN codes and ccs, with an ARI score of 0.89, closely followed by MLST (ARI = 0.55) (Figure 2) indicating that the ‘lineage’ threshold was concordant with LIN assignment and cc.

**Figure 2.**
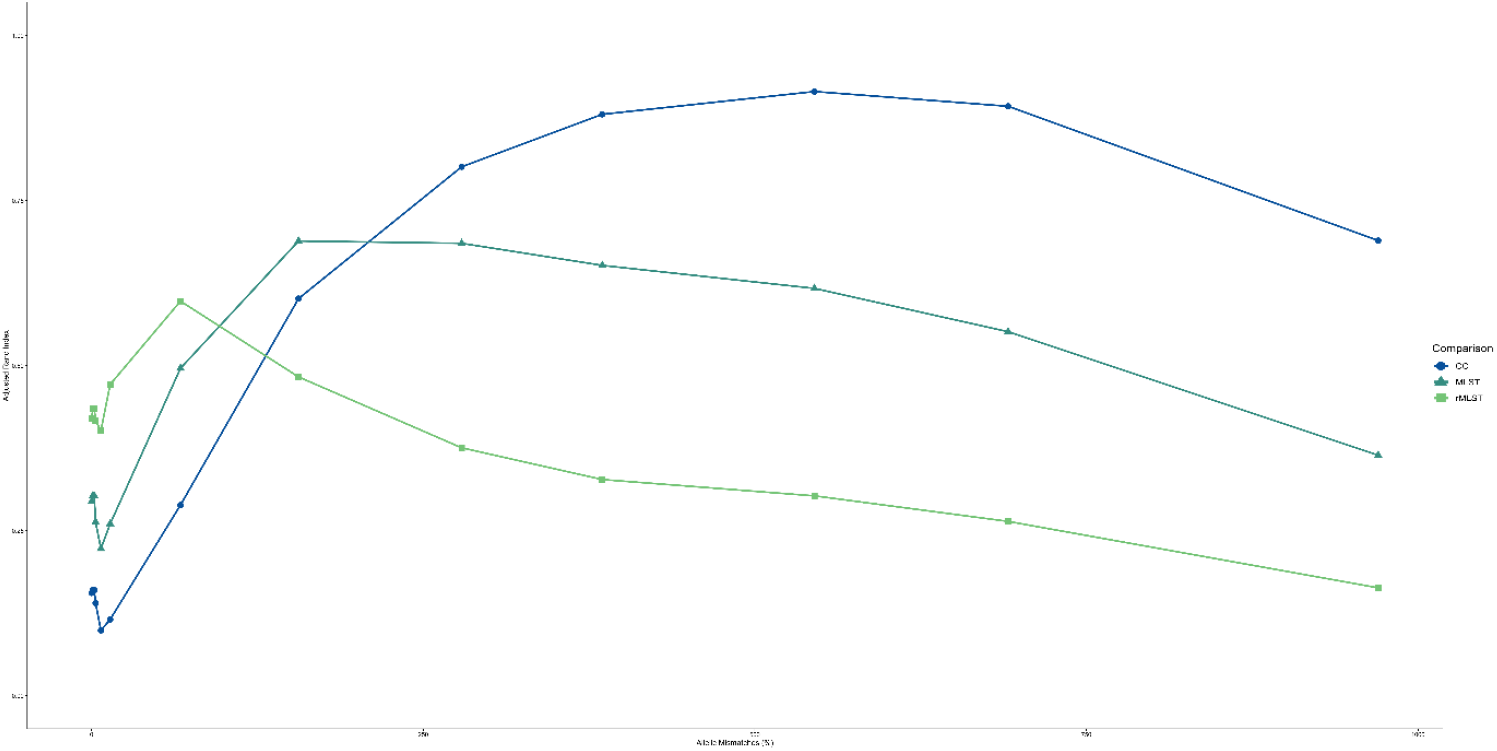
The adjusted Rand Index (aRandIndex) of the N. meningitidis allelic mismatch thresholds (%) (n = 6,131). Each line represents a different comparison metric: (i) clonal complex, (ii) core gene cluster 100, (iii) core gene cluster 200, (iv) ST, (v) rST.

### Implementation of the LIN thresholds

A total of 5,623 unique LIN codes were assigned in Dataset 1 (Supplementary Table 1). At the lineage threshold, 11 (12.4%) LIN codes were associated with more than one cc, therefore, 77.6% (38/49) of ccs had a unique lineage LIN assigned. Due to polyphyly within ccs, a cc may have different LIN assignments at the super-lineage and lineage threshold level. In consequence, overlapping assignment of some LIN codes at the higher thresholds occurred.

Dataset 2 (*n* = 1,888), consisting of 50 isolates from each cc, was used to identify concordance between 7-locus MLST profiles (ccs) and assigned LIN codes through phylogenetic analyses. When investigating 16 meningococcal ccs known to be associated with invasive disease, most were found to possess a representative LIN code at the lineage level (2^nd^ threshold). Instances where this differed were when the cc exhibited either sub-structure (cc269; 0_15 and 0_18) or inter-relationships (cc4 (57_1_0_2) and cc5 (57_1_0_0)) (Figure 3; Supplementary Table 2).

**Figure 3.**
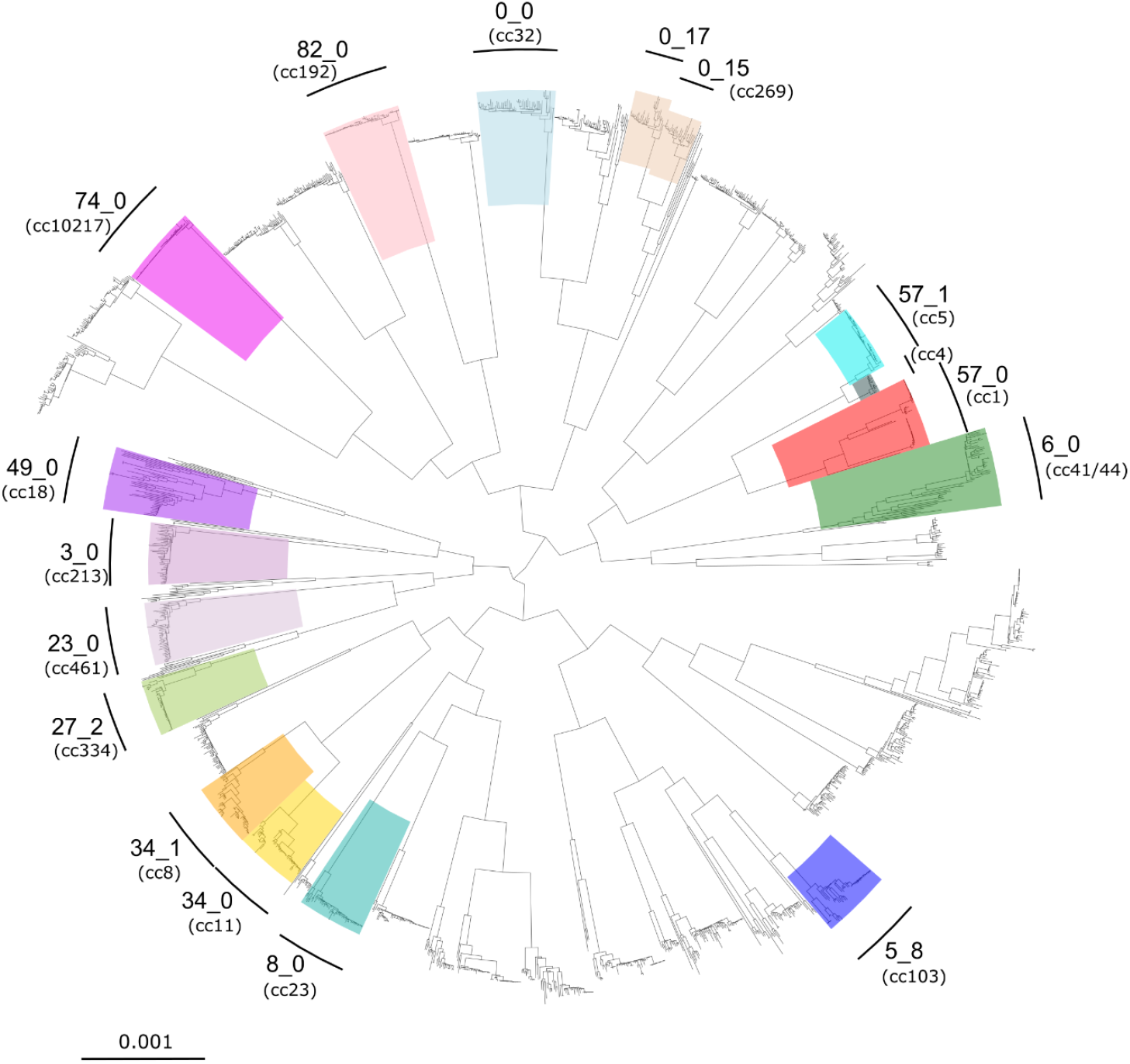
Maximum likelihood phylogeny of 1,888 N. meningitidis genomes, built using RAxML with 100 bootstraps and corrected for recombination using ClonalFrameML. The phylogenetic tree is representative of the meningococcal population, with up to 50 isolates selected from each clonal complex. Construction used the 1,329 cgMLST v3 loci. The 16 most relevant hyperinvasive clonal complexes are coloured, with their respective cgLINs annotated for the lineage level.

A total of 37,777 unique LIN codes were assigned to 62,330 distinct *N. meningitidis* cgSTs on PubMLST (12^th^ August 2025). LIN codes were assigned to 42,936/45,541 (94.28%) genomes. The remaining 2,605 (5.72%) isolates that did not have a LIN code lacked a cgST, which was likely due to inadequate genome quality. The unique LIN codes (*n* = 37,777 LIN codes) defined 118 super-lineages, 339 lineages, and 536 sub-lineages. Over half (56.26%) of the *N. meningitidis* genomes were represented by 3 groups (6, 34, and 0) at the super-lineage threshold. At the lineage threshold, there were two predominant LIN codes, corresponding to cc41/44 (6_0) and cc11 (34_0) (Figure 2), that accounted for 19.03% and 17.76% of the *N. meningitidis* genomes, respectively. At the highest resolution of 1 locus mismatch (12^th^ threshold), there were 37,702 clonal groups.

### Linking the LIN code thresholds to corresponding MLST lineages

For the 107 *N. meningitidis* reference isolates used to develop 7-locus MLST (Maiden *et al*., 1998), LIN codes were assigned to 91/107 (85.05%) isolates, all of which had a designated cgST (Supplementary Table 5). The ccs in this dataset were compared to the designated LIN thresholds (Table 2). A further two lineage designations for cc269 and one lineage designation for cc4240/6688 were identified, based on the LIN designations from Dataset 1. This represented the overall populations for each cc. The LIN codes at this 2^nd^ (lineage) threshold strongly corresponded to that of historical MLEE lineage, clonal complex, and gene-by-gene cgMLST lineage. The cc4 and cc5 capsule A isolates were not differentiated at the lineage threshold; however, they were resolved at the 4^th^ threshold (385 locus differences), with two unique LIN codes (Table 2; Supplementary Table 5).

**Table 2.**
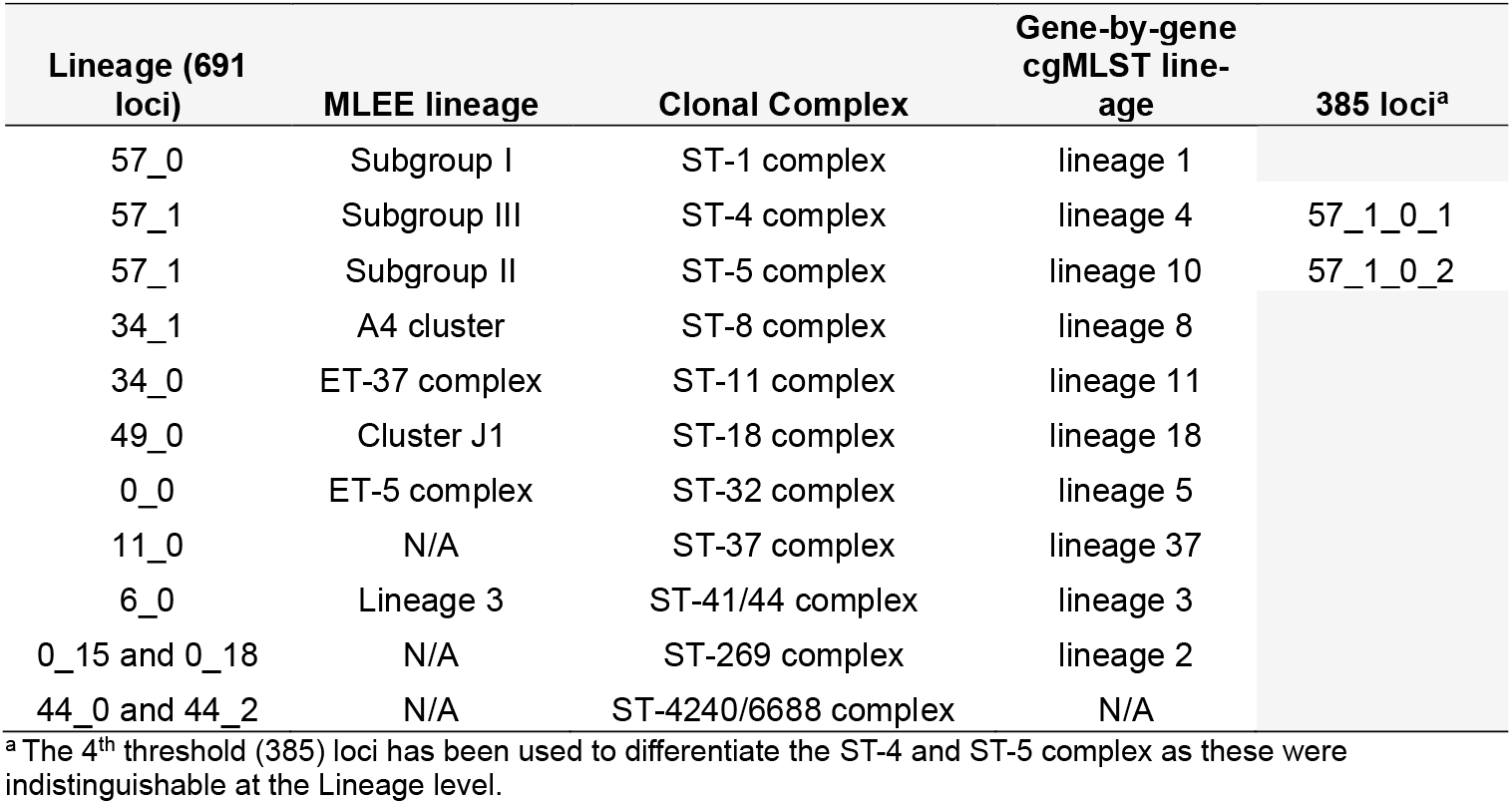
Backwards compatibility of LIN codes at the 2^nd^ threshold (“Lineage” level; 691 loci) compared to MLEE lineage, clonal complex, and cgMLST lineage for the top 11 hyperinvasive lineages.

### Application of LIN codes to published outbreaks

LIN codes were used to describe relationships among the genomes of 23 disease and carriage isolates from the 2011 meningococcal serogroup A cc5 (A:cc5) epidemic in Chad (Diallo *et al*., 2017). This resolved the three independent groups previously found by whole genome MLST (wgMLST) (Diallo *et al*., 2017), demonstrating that similar levels of discrimination were achieved using cgMLST or wgMLST based approaches (Maiden *et al*., 2013; Bratcher *et al*., 2014). At the 7-locus difference level (0.43% allelic mismatches) there were 6 unique LIN codes (Figure 4), with 13 isolates (*n* = 13; 56.5%) assigned to a single LIN, corresponding to cluster 2 in the published analysis (Diallo *et al*., 2017). The LIN codes discriminated cluster 2 into three groups at the 3-locus difference level (Figure 5). Two isolates within this cluster (PubMLST IDs 34997 and 34999) were indistinguishable, sharing the same 13-digit LIN code (Supplementary Table 6).

**Figure 4.**
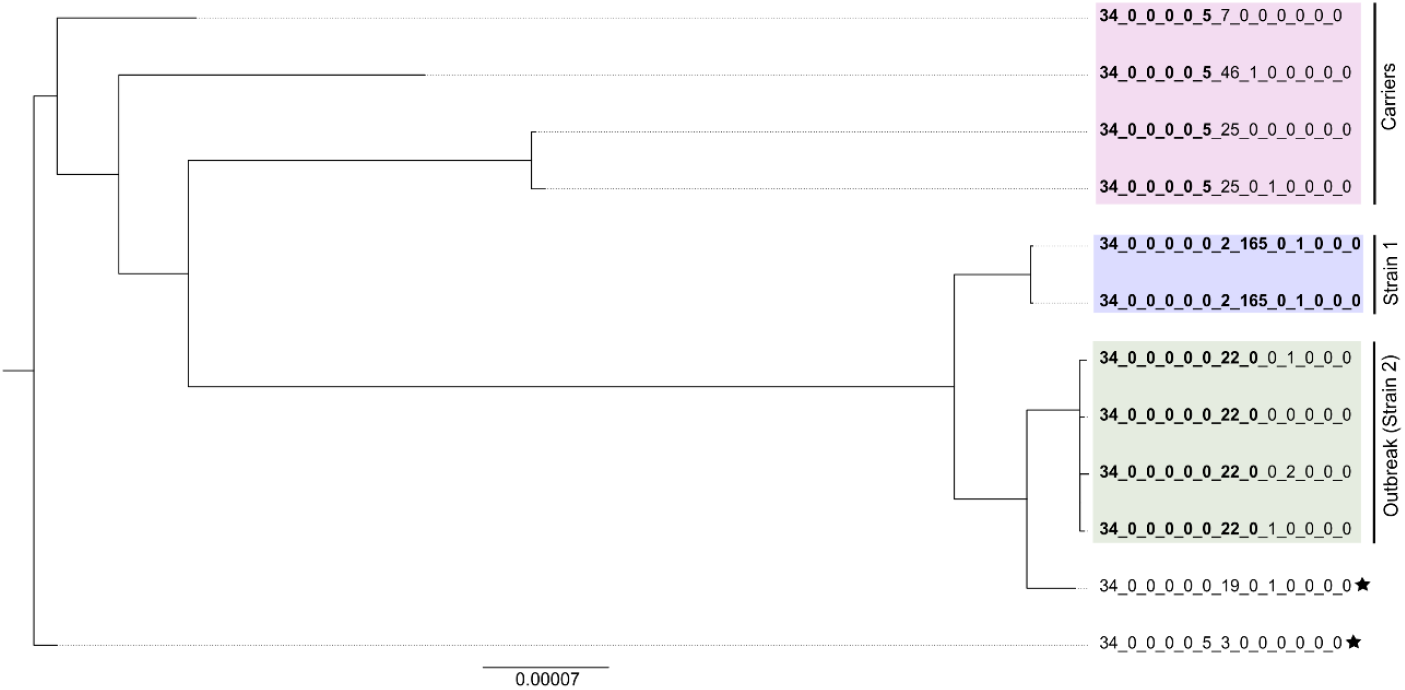
Maximum likelihood phylogeny of 12 N. meningitidis genomes, built using RAxML with 100 bootstraps and corrected for recombination using ClonalFrameML, which was previously analysed by Jolley *et al*. (2000). Construction used the 1,329 cgMLST v3 loci and has been rooted using genome ID 698 (FAM18) from PubMLST. Reference genomes are annotated with stars. Groupings of the isolates are annotated as per the initial analysis (i) Carriers, (ii) Strain 1, and (iii) Outbreak (Strain 2). The isolates have been annotated with their respective full cgLINs (to the 13th threshold). The bold lettering illustrates the similarities between the cgLINs within each grouping.

**Figure 5.**
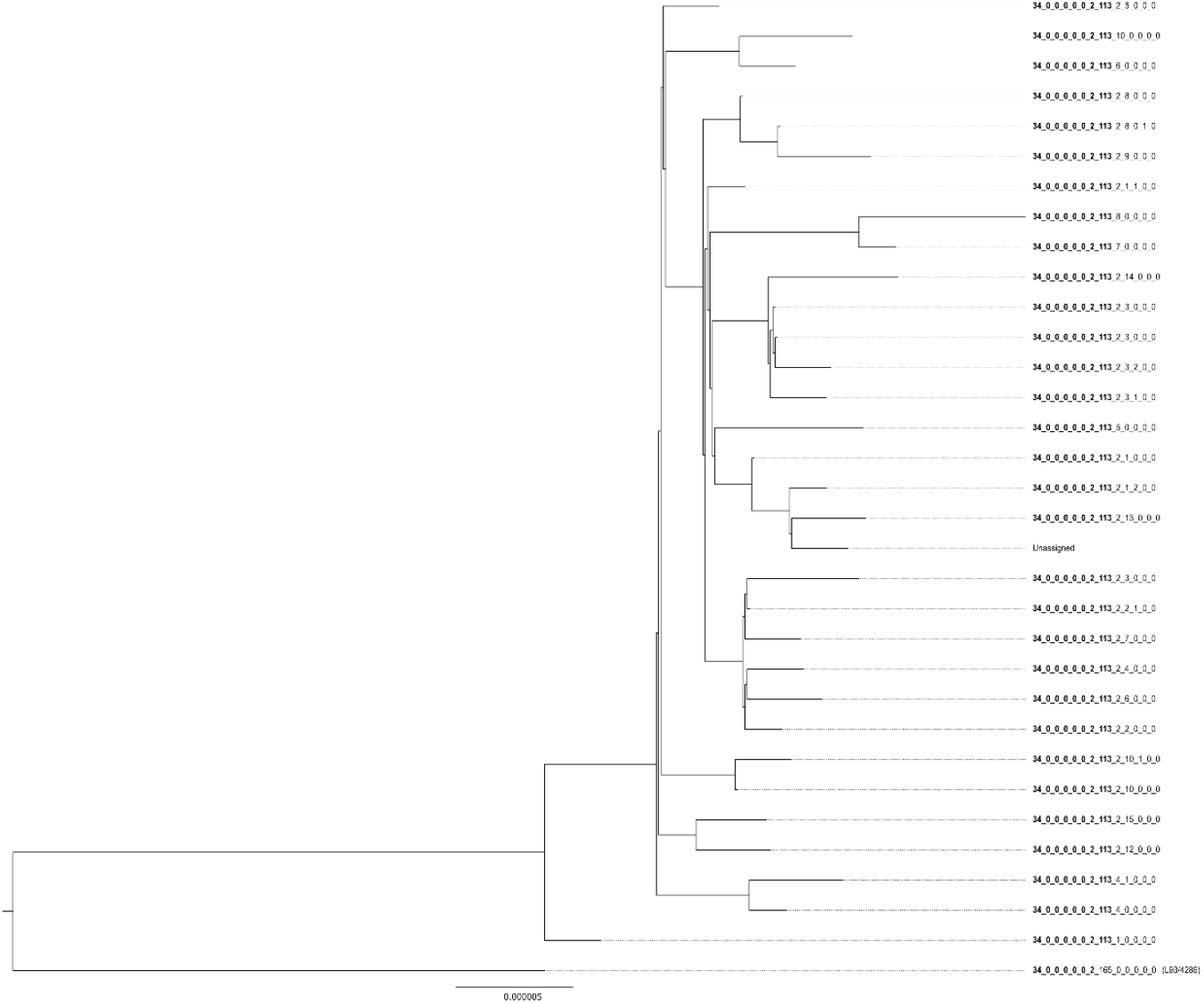
Maximum likelihood phylogeny of 33 N. meningitidis cc11:C genomes, built using RAxML with 100 bootstraps and corrected for recombination using ClonalFrameML, which was a subset from previous analysis by Jolley *et al*. (2012). Construction used the 1,329 cgMLST v3 loci and has been rooted using genome ID 644 (UK 1993 reference strain) from PubMLST. Reference genomes are annotated with stars. The isolates have been annotated with their respective full cgLINs (to the 13th threshold). Isolates that state ‘unassigned’ do not have an associated cgLIN. The bold lettering illustrates the similarities between the cgLINs.

LIN codes were also applied to the genomes of carriage and disease isolates obtained from a 1997 C:cc11 IMD outbreak at the University of Southampton in the United Kingdom (Gilmore *et al*., 1999; Jolley *et al*., 2012). The LIN codes accurately represented the relationships previously established by conventional MLST and antigen typing (Figure 5), but also further defined the outbreak cause as a distinct subvariant (defined by LIN code 34_0_0_0_0_0_22_0) of C:cc11 (defined by LIN code 34_0_0_0_0_0) that was causing cases of IMD globally in the 1990s. As of 6^th^ October 2025, there were 1,904 isolate genomes associated with this LIN code prefix recorded in the PubMLST *Neisseria spp*. (www.pubmlst.org/neisseria) database, which included both disease and carriage isolates obtained during a national outbreak in the Czech Republic in 1993 (Krízová *et al*., 1997; Jolley *et al*., 2000). The Czech C:cc11 isolates (*n* = 32) had indistinguishable LIN codes (sharing the LIN code prefix of 34_0_0_0_0_0) until the 7-locus mismatch threshold (Figure 6; Supplementary Table 7), which was unique within the PubMLST database (Jolley *et al*., 2018). There were no isolates from outside the Czech Republic (*n* = 44; Supplementary Table 7) that shared this LIN code at the 14-locus mismatch threshold (where the carriage isolates are indistinguishable). There were shared LIN codes among the Czech Republic carriage isolates and isolates from the C:cc11 outbreak in Southampton (Gilmore *et al*., 1999) at the 156 loci threshold (Supplementary Table 8), as well as other cc11 capsule B and C isolates from other countries and continents.

**Figure 6.**
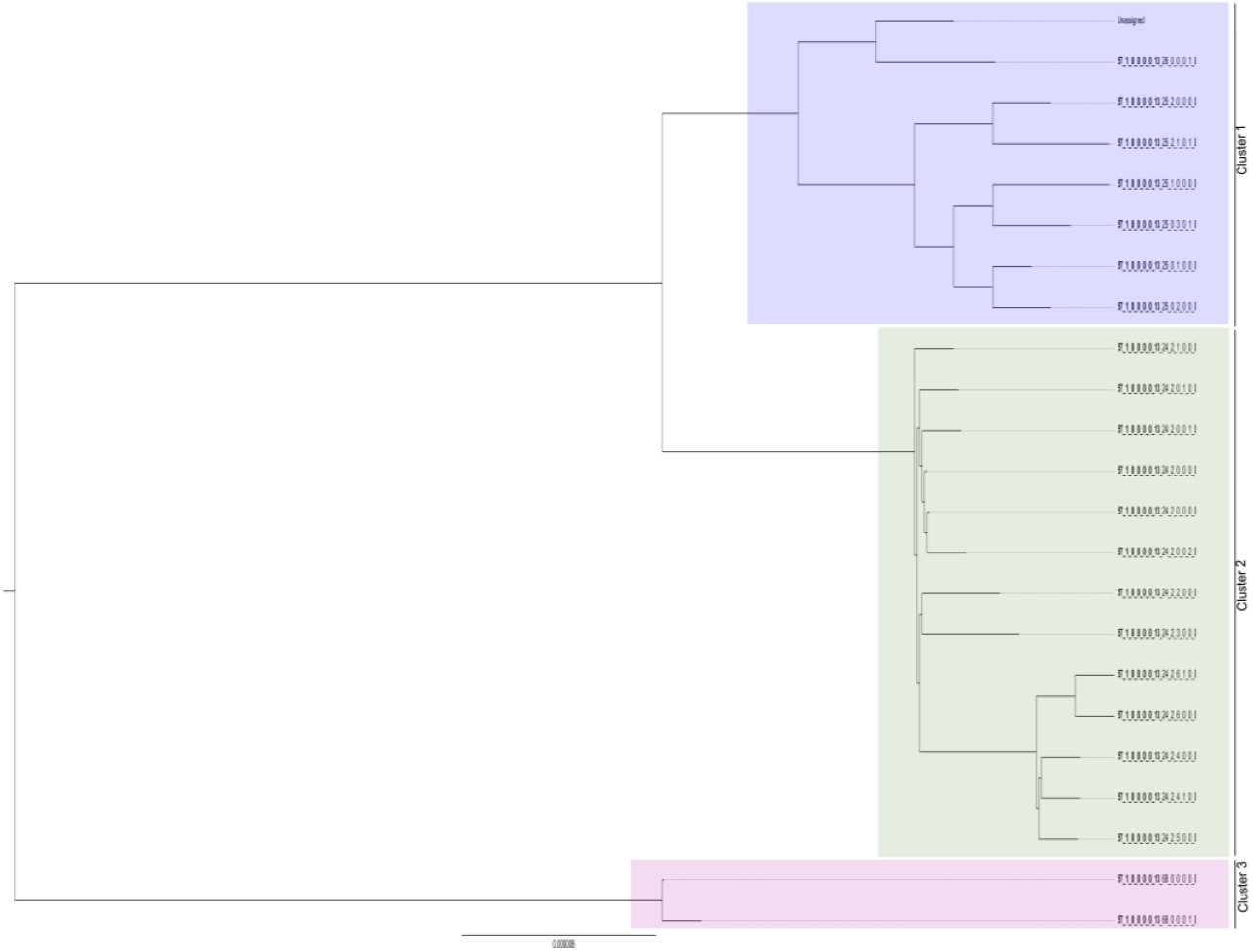
Maximum likelihood phylogeny of 23 N. meningitidis genomes, built using RAxML with 100 bootstraps and corrected for recombination using ClonalFrameML, which was previously analysed by Diallo *et al*. (2015). Construction used the 1,329 cgMLST v3 loci and has been rooted on the branch harbouring genome IDs 34995 and 34996 from PubMLST. Groupings of the isolates are annotated as per the initial analysis (i) Cluster 1, (ii) Cluster 2, and (iii) Cluster 3. The isolates have been annotated with their respective full cgLINs (to the 13th threshold). Isolates that state ‘unassigned’ do not have an associated cgLIN. The bold lettering illustrates the similarities between the cgLINs within each grouping.

## Discussion

Accurately describing and communicating bacterial diversity is essential for studies of population biology, pathology, epidemiology, and disease surveillance. Despite advances in the control of IMD though vaccination programmes, IMD remains a global public health threat (Vuocolo *et al*., 2018). Early detection and diagnosis are important for both the effective treatment and public health management, increasing patient survival and reducing long-term health consequences including disabling sequelae and mortality (Ciftci *et al*., 2025). Of particular concern at the time of writing was the emergence of novel hyperinvasive lineages, including some associated with antimicrobial resistance, such as the quinolone resistant cc4821 (Guo *et al*., 2018; Chen *et al*., 2021) and penicillin resistant cc23 meningococci (Potts *et al*., 2022).

From 1998 onwards, seven locus MLST and antigen fine typing was used very successfully to catalogue variants of *N. meningitidis*. This aided outbreak surveillance, defining the population structure, identifying emerging lineages, and assisting public health interventions such as mass vaccination campaigns (Maiden *et al*., 1998; Brehony *et al*., 2007; Jolley *et al*., 2007; Itsko *et al*., 2020). WGS technologies has permitted higher resolution analysis, including extended MLST schemes rMLST, wgMLST, and cgMLST (Maiden *et al*., 2013; Bratcher *et al*., 2014). Standardising these nomenclatures proved challenging as hundreds of loci were employed, resulting in thousands of unique STs. Here, a LIN code system was developed to meet this problem, through employing an updated cgMLST scheme (1,329 loci). The updated cgMLST v3 scheme was robust, automatable, and achieved high resolution with the resulting LIN code system providing a stable, scalable, 13-level typing system.

*N. meningitidis* is naturally transformable and exhibits extensive evidence of horizontal gene transfer (HGT) influencing its population structure and the evolution of variants (Schoen *et al*., 2009; Bratcher *et al*., 2012). Around 40% of the meningococcal core genes show evidence of recombination events caused by HGT (Joseph *et al*., 2011), particularly affecting genes that are involved in the pathogenic phenotype (Didelot and Maiden, 2010). While HGT can be a homogenising process, it also provides a mechanism for the generation and movement of genetic variation across unrelated lineages and amongst species (Bratcher *et al*., 2012). Examples of whole gene movement include the superoxide dismutase *sodC* gene, which appears to have been horizontally acquired from *Haemophilus influenzae* (Kroll *et al*., 1998), and the acquisition of *norB* and *aniA* from *N. gonorrhoeae*, that has resulted in a novel urogenital tract variant of cc11 (Tzeng *et al*., 2017; Tzeng *et al*., 2023).

There are two types of vaccines used to control IMD: (i) conjugate-polysaccharide vaccines, targeting serogroups A, C, W, Y, and X; and (ii) outer membrane vesicle (OMV) and protein-based vaccines, principally developed as substitutes to vaccines targeting serogroup B polysaccharide (Maiden and Stuart, 2002; Trotter and Maiden, 2009; Granoff, 2010; Vogel *et al*., 2013). The protein-based vaccines are susceptible to vaccine escape by variation in the targeted proteins, which can be generated by mutation or HGT (Mikucki and Kahler, 2023). This alters the relationship between genetic lineage, hyperinvasive phenotype and vaccine antigens. Application of the LIN code system facilitates the rapid and standardised assessment of these relationships, enhancing the capacity of public health practitioners to monitor and respond to changes in the variants responsible for IMD in their jurisdictions, including variation in vaccination policy.

Hyperinvasive meningococcal lineages persist for decades and cause endemic, hyper endemic epidemic and pandemic IMD (Caugant *et al*., 1986; Watkins and Maiden, 2012; Watkins and Maiden, 2017). They usually belong to one of six serogroups (A, B, C, W, X, and Y) that are associated with specific ccs, defined by MLST (Jolley *et al*., 2005; Mikucki and Kahler, 2023). These ccs are associated with particular LIN code prefixes, e.g. cc1, 57_0; cc4, 57_1_0_2; cc5, 57_1_0_0; cc8, 34_1; cc11, 34_0; cc32, 0_0; cc41/44, 6_0; cc103, 5_8; cc269, 0_17 and 0_15, cc334, 27_2; and cc461, 23_0 (Figure 3). Changes in carried hyperinvasive ccs, and their association with antigens, alter the *N. meningitidis* population structure in terms of IMD prevalence and nature (Jolley *et al*., 2005; Mikucki and Kahler, 2023). For example, serogroup A hyperinvasive meningococci (cc1, 57_0; cc4, 57_1_0_2; and cc5, 57_1_0_0) have been responsible for large-scale pandemic and epidemic IMD, particularly within the sub-Saharan African Meningitis belt (Olyhoek *et al*., 1987; Greenwood, 1999). In the earlier part of the 20^th^ century, A:cc1 dominated the region, being replaced by the emergence of A:cc5 associated with the Hajj pilgrimage in 1987. The A:cc5 meningococci underwent change over the subsequent 30-40 years. Variants of A:cc5 emerged during this time, including A:cc5:ST-7 and A:cc5:ST-2859 (differentiating at the 8^th^ LIN threshold; 14 loci mismatches), through introgression between different hyperinvasive lineages and ccs (Watkins and Maiden, 2017).

The epidemics of meningococcal serogroup A disease in the African meningitis belt were halted by the introduction of MenAfriVac in 2010 (Sow *et al*., 2011; Kristiansen *et al*., 2012; Maiden, 2013; Trotter *et al*., 2017). Meningococcal LIN codes were applied to clonal serogroup A isolates from a 2011 epidemic in Chad, collected during vaccine introduction, which were previously analysed using WGS techniques (Diallo *et al*., 2017). The LIN codes were able to replicate the clustering found in the original publication. Additionally, further information was provided by the LIN codes regarding indistinguishable isolates and further sub-divisions of the clusters that were not identified previously, showing that LIN codes enable investigation across different hierarchical levels of the meningococcal population.

PubMLST provides an easy-to-use interface, permitting the analysis and comparison of LIN codes among isolates through a “similar isolates” function and table on each isolate record page. This function identified that the C:cc11 (34_0) isolates recovered from the Southampton outbreak (Jolley *et al*., 2012) were members of the cc11 variant (34_0_0_0_0_0; 1,978 isolates in PubMLST on 21/12/2025) associated with epidemic disease in the 1990s. The outbreak isolates (C:cc11, ST-50) were unique (34_0_0_0_0_0_22; 4 isolates in PubMLST on 21^st^ December 2025), showing that this was likely a variant that emerged in the locale of Southampton, which has not been subsequently isolated. The isolates from the Czech Republic (Jolley *et al*., 2000) also belonged to the IMD C:cc11 in the 1990s (34_0_0_0_0_0). A subvariant of this group, found at the 67-locus difference threshold (34_0_0_0_0_0_2), has 553 isolates in PubMLST (accessed on 21^st^ December 2025) that have been isolated 1991-2024 from Europe, North America and South Africa. Of this, there were 44 isolates on PubMLST that are unique to the Czech Republic in the 1990s at the 14-locus difference threshold (34_0_0_0_0_0_2_113). This suggested that there was little to no spread of this lineage beyond the Czech Republic, and that this variant was likely introduced into the Czech Republic due to the increase of human mobility after the velvet revolution increased movement of and to the Czech population. This illustrates how LIN codes facilitate rapid and accurate identification of isolates belonging to hyperinvasive lineages, including those that contribute to outbreaks.

In conclusion, the LIN code approach presented here provides a stable high-resolution tool to catalogue, describe, and compare meningococcal variation. Full LIN codes are a computer readable system, with different LIN thresholds chosen to corroborate established nomenclatures (i.e. STs and ccs) to provide backwards compatibility and ease-of-use. The multi-hierarchal scheme distinguishes among closely related meningococcal variants and has application in epidemiological surveillance at all scales. This will aid studies of pathology and vaccine development, facilitating improvements in global public health through the identification and monitoring of emerging variants and lineages, including those of major concern such as those associated with vaccine escape or antibiotic resistance.

## References

Badur, S., M. Khalaf, S. Öztürk, R. Al-Raddadi, A. Amir, F. Farahat and A. Shibl (2022). “Meningococcal Disease and Immunization Activities in Hajj and Umrah Pilgrimage: a review.” Infect Dis Ther 11(4): 1343–1369.

Birtles, A., K. Hardy, S. J. Gray, S. Handford, E. B. Kaczmarski, V. Edwards-Jones and A. J. Fox (2005). “Multilocus sequence typing of Neisseria meningitidis directly from clinical samples and application of the method to the investigation of meningococcal disease case clusters.” J Clin Microbiol 43(12): 6007–6014.

Bratcher, H. B., J. S. Bennett and M. C. Maiden (2012). “Evolutionary and genomic insights into meningococcal biology.” Future Microbiol 7(7): 873–885.

Bratcher, H. B., C. Corton, K. A. Jolley, J. Parkhill and M. C. J. Maiden (2014). “A gene-by-gene population genomics platform: de novo assembly, annotation and genealogical analysis of 108 representative Neisseria meningitidis genomes.” BMC Genomics 15(1): 1138.

Brehony, C., K. A. Jolley and M. C. Maiden (2007). “Multilocus sequence typing for global surveillance of meningococcal disease.” FEMS Microbiol Rev 31(1): 15–26.

Cabellos, C., I. Pelegrín, E. Benavent, F. Gudiol, F. Tubau, D. Garcia-Somoza, R. Verdaguer, J. Ariza and P. Fernandez Viladrich (2019). “Invasive Meningococcal Disease: What We Should Know, Before It Comes Back.” Open Forum Infect Dis 6(3): ofz059.

Campbell, H., N. Andrews, S. R. Parikh, J. White, M. Edelstein, X. Bai, J. Lucidarme, R. Borrow, M. E. Ramsay and S. N. Ladhani (2022). “Impact of an adolescent meningococcal ACWY immunisation programme to control a national outbreak of group W meningococcal disease in England: a national surveillance and modelling study.” Lancet Child Adolesc Health 6(2): 96–105.

Caugant, D. A., L. O. Frøholm, K. Bøvre, E. Holten, C. E. Frasch, L. F. Mocca, W. D. Zollinger and R. K. Selander (1986). “Intercontinental spread of a genetically distinctive complex of clones of Neisseria meningitidis causing epidemic disease.” Proc Natl Acad Sci U S A 83(13): 4927–4931.

Caugant, D. A. and M. C. Maiden (2009). “Meningococcal carriage and disease--population biology and evolution.” Vaccine 27 Suppl 2(4): B64–70.

Caugant, D. A. and P. Nicolas (2007). “Molecular surveillance of meningococcal meningitis in Africa.” Vaccine 25: A8–A11.

Chen, M., O. B. Harrison, H. B. Bratcher, Z. Bo, K. A. Jolley, C. M. C. Rodrigues, J. E. Bray, Q. Guo, X. Zhang, M. Chen and M. C. J. Maiden (2021). “Evolution of Sequence Type 4821 Clonal Complex Hyperinvasive and Quinolone-Resistant Meningococci.” Emerg Infect Dis 27(4): 1110–1122.

Ciftci, E., D. Ocal, A. Somer, H. Tezer, D. Yilmaz, S. Bozkurt, O. U. Dursun, Ş. Merter and E. C. Dinleyici (2025). “Current methods in the diagnosis of invasive meningococcal disease.” Front Pediatr 13: 1511086.

Delgado-Blas, J. F., M. Rethoret-Pasty and S. Brisse (2025). “Life Identification Number (LIN) codes for the genomic taxonomy of Corynebacterium diphtheriae strains.” Genome Med 18(1): 5.

Diallo, K., K. Gamougam, D. M. Daugla, O. B. Harrison, J. E. Bray, D. A. Caugant, J. Lucidarme, C. L. Trotter, M. Hassan-King, J. M. Stuart, O. Manigart, B. M. Greenwood and M. C. J. Maiden (2017). “Hierarchical genomic analysis of carried and invasive serogroup A Neisseria meningitidis during the 2011 epidemic in Chad.” BMC Genomics 18(1): 398.

Didelot, X. and M. C. Maiden (2010). “Impact of recombination on bacterial evolution.” Trends Microbiol 18(7): 315–322.

Didelot, X. and D. J. Wilson (2015). “ClonalFrameML: Efficient Inference of Recombination in Whole Bacterial Genomes.” PLOS Computational Biology 11(2): e1004041.

Feavers, I. M., S. J. Gray, R. Urwin, J. E. Russell, J. A. Bygraves, E. B. Kaczmarski and M. C. Maiden (1999). “Multilocus sequence typing and antigen gene sequencing in the investigation of a meningococcal disease outbreak.” J Clin Microbiol 37(12): 3883–3887.

Gilmore, A., G. Jones, M. Barker, N. Soltanpoor and J. M. Stuart (1999). “Meningococcal disease at the University of Southampton: outbreak investigation.” Epidemiol Infect 123(2): 185–192.

Granoff, D. M. (2010). “Review of Meningococcal Group B Vaccines.” Clinical Infectious Diseases 50(Supplement_2): S54–S65.

Greenwood, B. (1999). “Manson lecture: meningococcal meningitis in Africa.” Transactions of the Royal Society of Tropical Medicine and Hygiene 93(4): 341–353.

Guo, Q., M. M. Mustapha, M. Chen, D. Qu, X. Zhang, M. Chen, Y. Doi, M. Wang and L. H. Harrison (2018). “Evolution of Sequence Type 4821 Clonal Complex Meningococcal Strains in China from Prequinolone to Quinolone Era, 1972-2013.” Emerg Infect Dis 24(4): 683–690.

Hahné, S. J., S. J. Gray, F. Jean, Aguilera, N. S. Crowcroft, T. Nichols, E. B. Kaczmarski and M. E. Ramsay (2002). “W135 meningococcal disease in England and Wales associated with Hajj 2000 and 2001.” Lancet 359(9306): 582–583.

Hamed, M. M., F. A. Mir, E. B. I. Elmagboul, A. Al-Khal, M. A. R. S. A. Maslamani, A. S. Deshmukh, H. E. Al-Romaihi, M. A. M. S. Janahi, F. B. Abid, A. S. A. Kashaf, G. Sher, V. K. Gupta, G. J. Wilson, J. Kadalayi and S. H. Doiphode (2021). “Molecular characteristics of Neisseria meningitidis in Qatar.” Scientific Reports 11(1): 4812.

Hennart, M., J. Guglielmini, S. Bridel, M. C. J. Maiden, K. A. Jolley, A. Criscuolo and S. Brisse (2022). “A Dual Barcoding Approach to Bacterial Strain Nomenclature: Genomic Taxonomy of Klebsiella pneumoniae Strains.” Mol Biol Evol 39(7).

Hubert, L. J. and P. Arabie (1985). “Comparing partitions.” Journal of Classification 2: 193–218.

Inkscape Team (2023). Inkscape, The Inkscape Project.

Itsko, M., A. C. Retchless, S. J. Joseph, A. Norris Turner, J. A. Bazan, A. Y. Sadji, R. Ouédraogo-Traoré and X. Wang (2020). “Full Molecular Typing of Neisseria meningitidis Directly from Clinical Specimens for Outbreak Investigation.” J Clin Microbiol 58(12).

Jansen van Rensburg, M. J., D. J. Berger, I. Yassine, D. Shaw, A. Fohrmann, J. E. Bray, K. A. Jolley, M. C. J. Maiden and A. B. Brueggemann (2024). “Development of the Pneumococcal Genome Library, a core genome multilocus sequence typing scheme, and a taxonomic life identification number barcoding system to investigate and define pneumococcal population structure.” Microb Genom 10(8).

Jolley, K. A., C. M. Bliss, J. S. Bennett, H. B. Bratcher, C. Brehony, F. M. Colles, H. Wimalarathna, O. B. Harrison, S. K. Sheppard, A. J. Cody and M. C. J. Maiden (2012). “Ribosomal multilocus sequence typing: universal characterization of bacteria from domain to strain.” Microbiology (Reading) 158(Pt 4): 1005–1015.

Jolley, K. A., J. E. Bray and M. C. J. Maiden (2018). “Open-access bacterial population genomics: BIGSdb software, the PubMLST.org website and their applications.” Wellcome Open Res 3: 124.

Jolley, K. A., C. Brehony and M. C. Maiden (2007). “Molecular typing of meningococci: recommendations for target choice and nomenclature.” FEMS Microbiol Rev 31(1): 89–96.

Jolley, K. A., D. M. Hill, H. B. Bratcher, O. B. Harrison, I. M. Feavers, J. Parkhill and M. C. Maiden (2012). “Resolution of a meningococcal disease outbreak from whole-genome sequence data with rapid Web-based analysis methods.” J Clin Microbiol 50(9): 3046–3053.

Jolley, K. A., J. Kalmusova, E. J. Feil, S. Gupta, M. Musilek, P. Kriz and M. C. Maiden (2000). “Carried meningococci in the Czech Republic: a diverse recombining population.” J Clin Microbiol 38(12): 4492–4498.

Jolley, K. A., D. J. Wilson, P. Kriz, G. McVean and M. C. Maiden (2005). “The influence of mutation, recombination, population history, and selection on patterns of genetic diversity in Neisseria meningitidis.” Mol Biol Evol 22(3): 562–569.

Joseph, B., R. F. Schwarz, B. Linke, J. Blom, A. Becker, H. Claus, A. Goesmann, M. Frosch, T. Müller, U. Vogel and C. Schoen (2011). “Virulence Evolution of the Human Pathogen Neisseria meningitidis by Recombination in the Core and Accessory Genome.” PLOS ONE 6(4): e18441.

Kristiansen, P. A., F. Diomandé, A. K. Ba, I. Sanou, A. S. Ouédraogo, R. Ouédraogo, L. Sangaré, D. Kandolo, F. Aké, I. M. Saga, T. A. Clark, L. Misegades, S. W. Martin, J. D. Thomas, S. R. Tiendrebeogo, M. Hassan-King, M. H. Djingarey, N. E. Messonnier, M.-P. Préziosi, F. M. LaForce and D. A. Caugant (2012). “Impact of the Serogroup A Meningococcal Conjugate Vaccine, MenAfriVac, on Carriage and Herd Immunity.” Clinical Infectious Diseases 56(3): 354–363.

Krízová, P., M. Musílek and J. Kalmusová (1997). “Development of the epidemiological situation in invasive meningococcal disease in the Czech Republic caused by emerging Neisseria meningitidis clone ET-15/37.” Cent Eur J Public Health 5(4): 214–218.

Kroll, J. S., K. E. Wilks, J. L. Farrant and P. R. Langford (1998). “Natural genetic exchange between Haemophilus and Neisseria: intergeneric transfer of chromosomal genes between major human pathogens.” Proc Natl Acad Sci U S A 95(21): 12381–12385.

Lingappa, J. R., A. M. Al-Rabeah, R. Hajjeh, T. Mustafa, A. Fatani, T. Al-Bassam, A. Badukhan, A. Turkistani, S. Makki, N. Al-Hamdan, M. Al-Jeffri, Y. Al Mazrou, B. A. Perkins, T. Popovic, L. W. Mayer and N. E. Rosenstein (2003). “Serogroup W-135 meningococcal disease during the Hajj, 2000.” Emerg Infect Dis 9(6): 665–671.

Lucidarme, J., D. M. C. Hill, H. B. Bratcher, S. J. Gray, M. du Plessis, R. S. W. Tsang, J. A. Vazquez, M.-K. Taha, M. Ceyhan, A. M. Efron, M. C. Gorla, J. Findlow, K. A. Jolley, M. C. J. Maiden and R. Borrow (2015). “Genomic resolution of an aggressive, widespread, diverse and expanding meningococcal serogroup B, C and W lineage.” Journal of Infection 71(5): 544–552.

Maiden, M. C. (2013). “The endgame for serogroup a meningococcal disease in Africa?” Clin Infect Dis 56(3): 364–366.

Maiden, M. C., J. A. Bygraves, E. Feil, G. Morelli, J. E. Russell, R. Urwin, Q. Zhang, J. Zhou, K. Zurth, D. A. Caugant, I. M. Feavers, M. Achtman and B. G. Spratt (1998). “Multilocus sequence typing: a portable approach to the identification of clones within populations of pathogenic microorganisms.” Proc Natl Acad Sci U S A 95(6): 3140–3145.

Maiden, M. C., M. J. Jansen van Rensburg, J. E. Bray, S. G. Earle, S. A. Ford, K. A. Jolley and N. D. McCarthy (2013). “MLST revisited: the gene-by-gene approach to bacterial genomics.” Nat Rev Microbiol 11(10): 728–736.

Maiden, M. C. and J. M. Stuart (2002). “Carriage of serogroup C meningococci 1 year after meningococcal C conjugate polysaccharide vaccination.” Lancet 359(9320): 1829–1831.

Mbaeyi, S. A., C. H. Bozio, J. Duffy, L. G. Rubin, S. Hariri, D. S. Stephens and J. R. MacNeil (2020). “Meningococcal Vaccination: Recommendations of the Advisory Committee on Immunization Practices, United States, 2020.” MMWR Recomm Rep 69(9): 1–41.

Mikucki, A. and C. M. Kahler (2023). “Microevolution and Its Impact on Hypervirulence, Antimicrobial Resistance, and Vaccine Escape in Neisseria meningitidis.” Microorganisms 11(12).

Olyhoek, T., B. A. Crowe and M. Achtman (1987). “Clonal population structure of Neisseria meningitidis serogroup A isolated from epidemics and pandemics between 1915 and 1983.” Reviews of infectious diseases 9(4): 665–692.

Palma, F., M. Hennart, K. A. Jolley, C. Crestani, K. L. Wyres, S. Bridel, C. A. Yeats, B. Brancotte, B. Raffestin, S. David, M. M. C. Lam, R. Izdebski, V. Passet, C. Rodrigues, M. Rethoret-Pasty, M. C. J. Maiden, D. M. Aanensen, K. E. Holt, A. Criscuolo and S. Brisse (2024). “Bacterial strain nomenclature in the genomic era: Life Identification Numbers using a gene-by-gene approach.” bioRxiv: 2024.2003.2011.584534.

Pardo de Santayana, C., M. Tin Tin Htar, J. Findlow and P. Balmer (2023). “Epidemiology of invasive meningococcal disease worldwide from 2010-2019: a literature review.” Epidemiol Infect 151: e57.

Potts, C. C., L. D. Rodriguez-Rivera, A. C. Retchless, F. Hu, H. Marjuki, A. E. Blain, L. A. McNamara and X. Wang (2022). “Antimicrobial Susceptibility Survey of Invasive Neisseria meningitidis, United States 2012–2016.” The Journal of Infectious Diseases 225(11): 1871–1875.

Rambaut, A. (2018). Figtree. Institute of Evolutionary Biology, University of Edinburgh, Edinburgh.

Reeves, M. W., B. A. Perkins and J. D. Wenger (1995). “Epidemic-associated Neisseria meningitidis detected by multilocus enzyme electrophoresis.” Emerg Infect Dis 1(2): 53–54.

Retchless, A. C., A. Chen, H. Y. Chang, A. E. Blain, L. A. McNamara, M. M. Mustapha, L. H. Harrison and X. Wang (2021). “Using Neisseria meningitidis genomic diversity to inform outbreak strain identification.” PLoS Pathog 17(5): e1009586.

RStudio Team (2020). RStudio: Integrated Development for R. RStudio. PBC, Boston, MA.

Schoen, C., H. Tettelin, J. Parkhill and M. Frosch (2009). “Genome flexibility in Neisseria meningitidis.” Vaccine 27 Suppl 2(Suppl 2): B103–111.

Scrucca, L., C. Fraley, T. Murphy and A. Raftery (2023). Model-Based Clustering, Classification, and Density Estimation Using mclust in R. New York, Chapman and Hall/CRC.

Slavinska, A., M. Kowalczyk, A. Kirkliauskienė, G. Vizuje, P. Siedlecki, J. Bikulčienė, K. Tamošiūnienė, A. Petrutienė and N. Kuisiene (2024). “Genetic characterization of Neisseria meningitidis isolates recovered from patients with invasive meningococcal disease in Lithuania.” Front Cell Infect Microbiol 14:p1432197.

Sow, S. O., B. J. Okoko, A. Diallo, S. Viviani, R. Borrow, G. Carlone, M. Tapia, A. K. Akinsola, P. Arduin, H. Findlow, C. Elie, F. C. Haidara, R. A. Adegbola, D. Diop, V. Parulekar, J. Chaumont, L. Martellet, F. Diallo, O. T. Idoko, Y. Tang, B. D. Plikaytis, P. S. Kulkarni, E. Marchetti, F. M. LaForce and M. P. Preziosi (2011). “Immunogenicity and safety of a meningococcal A conjugate vaccine in Africans.” N Engl J Med 364(24): 2293–2304.

Stamatakis, A. (2006). “RAxML-VI-HPC: maximum likelihood-based phylogenetic analyses with thousands of taxa and mixed models.” Bioinformatics 22(21): 2688–2690.

Stephens, D. S., B. Greenwood and P. Brandtzaeg (2007). “Epidemic meningitis, meningococcaemia, and Neisseria meningitidis.” Lancet 369(9580): 2196–2210.

Stollenwerk, N., M. C. Maiden and V. A. Jansen (2004). “Diversity in pathogenicity can cause outbreaks of meningococcal disease.” Proc Natl Acad Sci U S A 101(27): 10229–10234.

Taha, M.-K., M. Achtman, J.-M. Alonso, B. Greenwood, M. Ramsay, A. Fox, S. Gray and E. Kaczmarski (2000). “Serogroup W135 meningococcal disease in Hajj pilgrims.” The Lancet 356(9248): 2159.

Trotter, C. L., C. Lingani, K. Fernandez, L. V. Cooper, A. Bita, C. Tevi-Benissan, O. Ronveaux, M.-P. Préziosi and J. M. Stuart (2017). “Impact of MenAfriVac in nine countries of the African meningitis belt, 2010–15: an analysis of surveillance data.” The Lancet Infectious Diseases 17(8): 867–872.

Trotter, C. L. and M. C. Maiden (2009). “Meningococcal vaccines and herd immunity: lessons learned from serogroup C conjugate vaccination programs.” Expert Rev Vaccines 8(7): 851–861.

Tzeng, Y. L., J. A. Bazan, A. N. Turner, X. Wang, A. C. Retchless, T. D. Read, E. Toh, D. E. Nelson, C. Del Rio and D. S. Stephens (2017). “Emergence of a new Neisseria meningitidis clonal complex 11 lineage 11.2 clade as an effective urogenital pathogen.” Proc Natl Acad Sci U S A 114(16): 4237–4242.

Tzeng, Y. L., S. Sannigrahi, Z. Berman, E. Bourne, J. L. Edwards, J. A. Bazan, A. N. Turner, J. W. B. Moir and D. S. Stephens (2023). “Acquisition of Gonococcal AniA-NorB Pathway by the Neisseria meningitidis Urethritis Clade Confers Denitrifying and Microaerobic Respiration Advantages for Urogenital Adaptation.” Infect Immun 91(5): e0007923.

Tzeng, Y. L. and D. S. Stephens (2021). “A Narrative Review of the W, X, Y, E, and NG of Meningococcal Disease: Emerging Capsular Groups, Pathotypes, and Global Control.” Microorganisms 9(3).

Unitt, A., M. Krisna, K. M. Parfitt, K. A. Jolley, M. C. J. Maiden and O. B. Harrison (2025). Neisseria gonorrhoeae LIN codes: a Robust, Multi-Resolution Lineage Nomenclature, eLife Sciences Publications, Ltd.

Vinatzer, B. A., A. J. Weisberg, C. L. Monteil, H. A. Elmarakeby, S. K. Sheppard and L. S. Heath (2017). “A Proposal for a Genome Similarity-Based Taxonomy for Plant-Pathogenic Bacteria that Is Sufficiently Precise to Reflect Phylogeny, Host Range, and Outbreak Affiliation Applied to Pseudomonas syringae sensu lato as a Proof of Concept.” Phytopathology 107(1): 18–28.

Vogel, U., M. K. Taha, J. A. Vazquez, J. Findlow, H. Claus, P. Stefanelli, D. A. Caugant, P. Kriz, R. Abad, S. Bambini, A. Carannante, A. E. Deghmane, C. Fazio, M. Frosch, G. Frosi, S. Gilchrist, M. M. Giuliani, E. Hong, M. Ledroit, P. G. Lovaglio, J. Lucidarme, M. Musilek, A. Muzzi, J. Oksnes, F. Rigat, L. Orlandi, M. Stella, D. Thompson, M. Pizza, R. Rappuoli, D. Serruto, M. Comanducci, G. Boccadifuoco, J. J. Donnelly, D. Medini and R. Borrow (2013). “Predicted strain coverage of a meningococcal multicomponent vaccine (4CMenB) in Europe: a qualitative and quantitative assessment.” Lancet Infect Dis 13(5): 416–425.

Vuocolo, S., P. Balmer, W. C. Gruber, K. U. Jansen, A. S. Anderson, J. L. Perez and L. J. York (2018). “Vaccination strategies for the prevention of meningococcal disease.” Hum Vaccin Immunother 14(5): 1203–1215.

Watkins, E. R. and M. C. Maiden (2017). “Metabolic shift in the emergence of hyperinvasive pandemic meningococcal lineages.” Sci Rep 7: 41126.

Watkins, E. R. and M. C. J. Maiden (2012). “Persistence of Hyperinvasive Meningococcal Strain Types during Global Spread as Recorded in the PubMLST Database.” PLOS ONE 7(9): e45349.

Wickham, H. (2007). “Reshaping Data with the reshape Package.” Journal of Statistical Software 21: 1–20.

Wickham, H. (2016). ggplot2: Elegant Graphics for Data Analysis. Springer-Verlag New York.

Yu, G., D. K. Smith, H. Zhu, Y. Guan and T. T.-Y. Lam (2017). “ggtree: an r package for visualization and annotation of phylogenetic trees with their covariates and other associated data.” Methods in Ecology and Evolution 8(1): 28–36.

