## Supplementary figures and images for "A Life Identification Number Barcoding (LIN Code) System for *Neisseria meningitidis*: high resolution multi-level typing of meningococci"

### Supplementary Figure 1

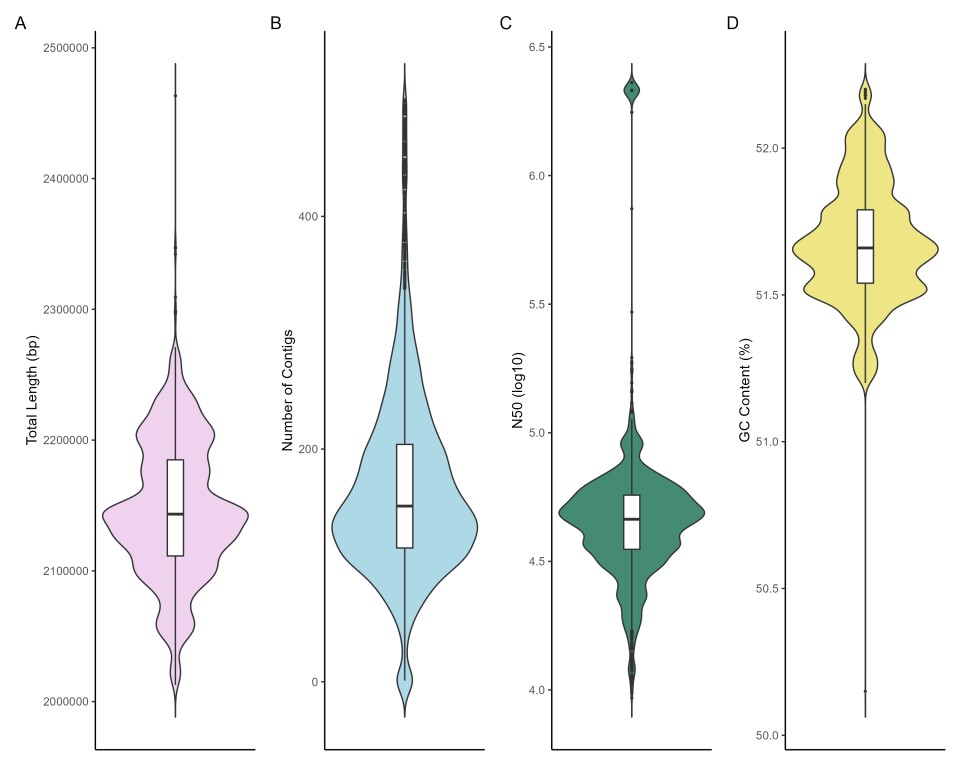
